# Evolution and Engineering of Allosteric Regulation in Protein Kinases

**DOI:** 10.1101/189761

**Authors:** David Pincus, Jai P. Pandey, Pau Creixell, Orna Resnekov, Kimberly A. Reynolds

## Abstract

Allosteric regulation – the control of protein function by sites far from the active site – is a common feature of proteins that enables dynamic cellular responses. Reversible modifications such as phosphorylation are well suited to mediate such regulatory dynamics, yet the evolution of new allosteric regulation demands explanation. To understand this, we mutationally scanned the surface of a prototypical kinase to identify readily evolvable phosphorylation sites. The data reveal a set of spatially distributed “hotspots” that coevolve with the active site and preferentially modulate kinase activity. By engineering simple consensus phosphorylation sites at these hotspots we successfully rewired *in vivo* cell signaling. Beyond synthetic biology, the hotspots are frequently used by the diversity of natural allosteric regulatory mechanisms in the kinase family and exploited in human disease.

**ONE SENTENCE SUMMARY:** Cell signaling is easily rewired by introducing new phosphoregulation at latent allosteric surface sites.

## MAIN TEXT

Allosteric regulation requires the cooperative action of many amino acids to functionally link distantly positioned amino acids. As a consequence, it is difficult to understand how allostery can evolve through a process of stepwise variation and selection. However, members of a protein family often display diverse regulatory mechanisms, suggesting that despite the complex intramolecular cooperativity required, allostery evolves readily (*1*). A potential explanation for how this might occur comes from separate lines of work that indicate a latent capacity for regulation at a diversity of surfaces in proteins. For example, it is possible to engineer synthetic allosteric switches through domain insertion at certain surface sites (*2–6*), and screens for small molecules that modify protein function sometimes identify cryptic allosteric regulatory sites (*7, 8*). In addition, experimental analysis of regulation in orthologs of the yeast MAP kinase Fus3 indicates that the capacity for allosteric regulation existed well before the regulatory mechanism evolved (*9*). Taken together, these findings suggest that proteins have an internal architecture in which a few sites on the protein surface are functionally “pre-wired” to provide control of protein active sites (*10*). This pre-wiring has been proposed to result not as a consequence of the need for regulation, but simply from the need for proteins to be evolvable (*11*). Thus, the acquisition of new regulation might amount to engaging or activating preexisting allosteric networks, a route to the evolution of regulation that is consistent with stepwise variation and selection.

An excellent model to test this proposal is the eukaryotic protein kinases (EPKs), a protein family that has diversified to control a vast array of cellular signaling activities. The EPKs catalyze the transfer of a phosphate group from adenosine triphosphate (ATP) onto a Ser/Thr/Tyr residue of a substrate protein, a reaction that is subject to regulation by different mechanisms at many surface regions in members of the kinase family (Fig. 1), including proteinprotein interactions, auto-inhibition, dimerization, and post-translational modification (*12*). Recently, Ferrell and colleagues proposed an idea for the evolution of one such mechanism – phosphoregulation – in which phosphorylation of a Ser/Thr/Tyr surface residue regulates protein activity. Since phosphorylation introduces a negative charge, the idea is that phosphoregulation might evolve simply by mutating an allosterically pre-coupled negatively charged residue (Asp/Glu) to a phosphorylatable residue (Ser/Thr/Tyr) (*13*). Thus, a constitutive negative charge at a latent allosteric site can be transformed into a regulated negative charge in a potentially stepwise manner (*14*).

**Figure 1.**
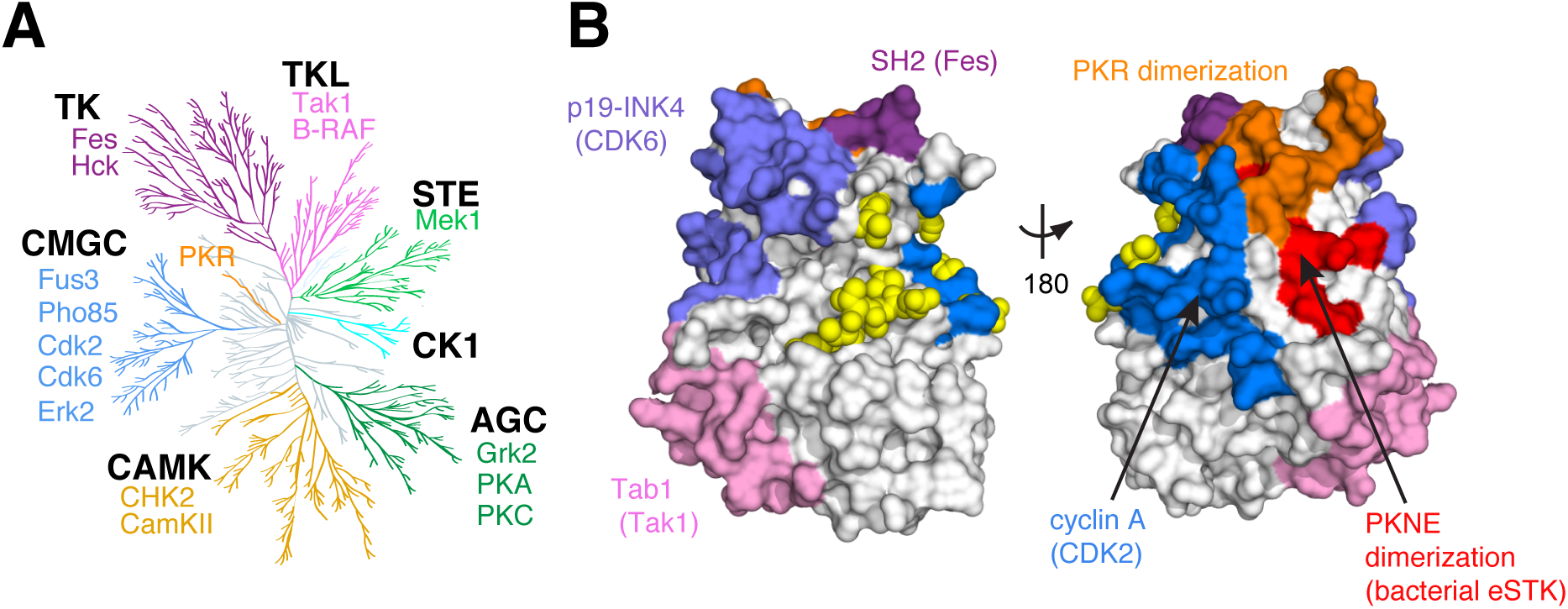
Regulatory Diversity in the Eukaryotic Protein Kinases. **A.** Unanchored dendrogram of the human kinome illustrating the diversity of the EPK superfamily and subfamilies. Individual subfamily members with functional mutations shown in Fig. 4c and included in Supplementary Table 7 are listed. TK: tyrosine kinase; TKL: TK-like; STE: STE7/11/20; CK1: Casein Kinase 1; AGC: protein kinase A/G/C; CAMK: Calmodulin kinase; CMGC: cyclin dependent kinase (CDK)/mitogen activated protein kinase (MAPK)/glycogen synthase kinase (GSK)/CDK-like kinase (CLK). **B.** Allosteric regulatory sites from diverse kinases mapped to a single representative structure - yeast CDK Pho85 (PDB: 2PK9, shown as space-filled surface). Regulatory surfaces were identified by structural alignment of the kinase of interest to Pho85; all Pho85 positions within 4Å of the interaction surface are colored. Color coding is the same as in (**A**). This mapping shows that regulation occurs at structurally diverse sites across the kinase structure.

To experimentally test this idea, we used the prototypical yeast CMGC kinase Kss1 as a model (see Supplementary Text for extended Kss1 background). Kss1 is a homolog of human ERK and is involved in signal transduction pathways that regulate yeast filamentous growth and the mating response (*15-18*). Kss1 activity can be quantitatively monitored in living yeast cells by its ability to specifically activate fluorescent transcriptional reporters of the mating pheromone response in the absence of its paralog, Fus3 (Fig. 2A). We conducted an unbiased alanine scan of all 40 Asp/Glu residues on the surface of Kss1 to determine which positions are functionally coupled to kinase activity. We integrated the resulting 40 Kss1 mutants as the only copy of Kss1 in the yeast genome, tagged at their C-terminus with a 3×FLAG epitope (Supplementary Tables 1and 2). To test their activity, we assayed for induction of the pheromone-responsive *AGA1pr-* YFP reporter at four concentrations of the alpha factor mating pheromone (αF) by flow cytometry. Though all mutants maintained wild type-like expression levels, nine mutations altered *in vivo* kinase activity (Fig. 2B, C, Supplementary Fig. 1A). Three of these positions were identified as Kss1 mutants with a functional effect in previous studies (D117, D156, D321, Supplementary Table 3) (*19, 20*). Though enriched in the N-terminal half of the primary Kss1 sequence, these nine mutations occur at positions distributed broadly over the Kss1 atomic structure - consistent with the notion that certain surface sites are selectively pre-wired to allosterically influence active site function.

**Figure 2.**
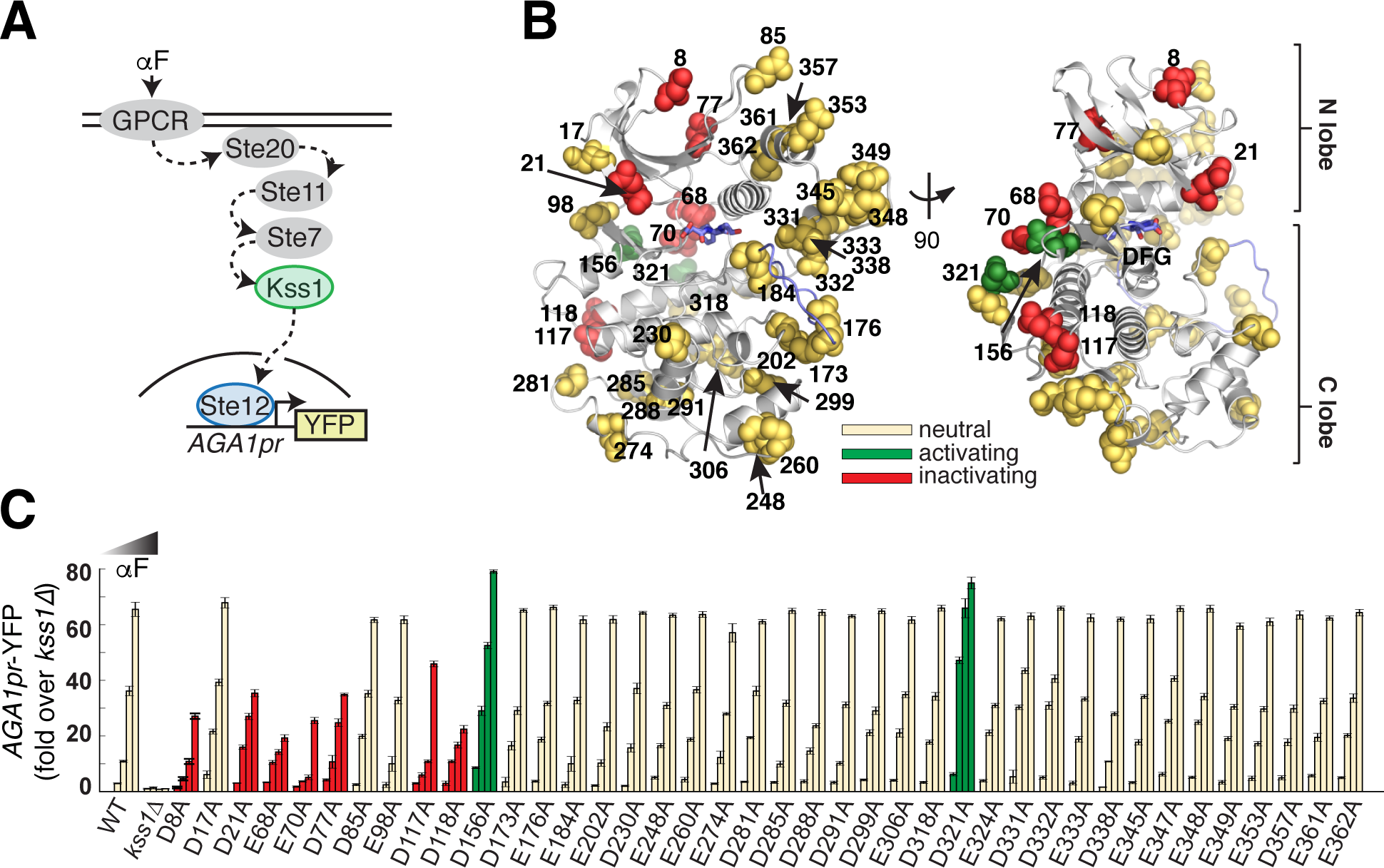
Alanine scan of acidic residues on the solvent accessible surface of yeast MAPK Kss1. **A.** Schematic of the Kss1-dependent yeast pheromone pathway. The alpha factor (αF) mating pheromone binds to a G-protein coupled receptor (GPCR), leading to activation of a signaling cascade culminating at the MAPK Kss1. Kss1 then activates the Ste12 transcription factor to induce the mating transcriptional program, which can be monitored by fusing the promoter of the target gene *AGA1* to a YFP reporter. **B.** Ribbon diagram of a Kss1 homology model (34) with the 40 solvent accessible Asp/Glu residues shown as spheres. The DFG motif and activation loop are indicated in light blue. All 40 positions were mutated individually to alanine to remove negative charge. **C.** The 40 resulting yeast strains along with WT and *kss1*Δ controls were assayed for activation of the *AGA1pr-YFP* reporter by flow cytometry following treatment with 0, 0.01, 0.1 and 1 μM αF for 4 hours. Bars represent the average of the median YFP fluorescence from 3 biological replicates normalized to the untreated *kss1*Δ cells, and error bars are the standard deviation of the biological replicates. Mutations at red and green positions resulted in significantly reduced or increased YFP expression (*p* < 0.05) in response to at least two doses of aF, respectively. Yellow positions indicate that the mutation had no effect in this assay. The color coding is identical in (**B**). The data show that nine acidic positions on the solvent accessible surface are functionally coupled to kinase activity.

We next tested whether these positions can support new regulation of Kss1 through phosphorylation by another yeast kinase *in vivo.* In principle, this gain of function can effectively rewire signaling through the mating response pathway. We chose to engineer regulation of Kss1 by protein kinase A (PKA) because the PKA substrate consensus motif, RRxS/T requires minimal local modifications, its activity in yeast cells is orthogonal to the pheromone pathway, and it can be hyper-activated in yeast via ectopic expression of Ras2^G19V^ (Fig. 3A, Supplementary Fig. 1B, C). We selected three of the nine mutationally sensitive positions (D8, E68 and E70) with the highest PKA substrate scores predicted by the computational tool pkaPS (1). To claim PKA-mediated allosteric regulation of Kss1, we must demonstrate: 1) that Kss1 retains functionality following introduction of a local PKA consensus motif (RRx**D/E**, termed pka-D/E); 2) that Kss1 loses activity when the charge is neutralized (RRx**A**, termed pka-A); and 3) that Kss1 now displays PKA-dependent activity *in vivo* with introduction of a phosphorylatable residue (RRx**S**, termed pka-S) (Supplementary Fig. 1d). In this manner, a functional surface negative charged residue can neutrally acquire a substrate consensus sequence for a kinase and become a phosphoregulatory site with one step of variation.

**Figure 3.**
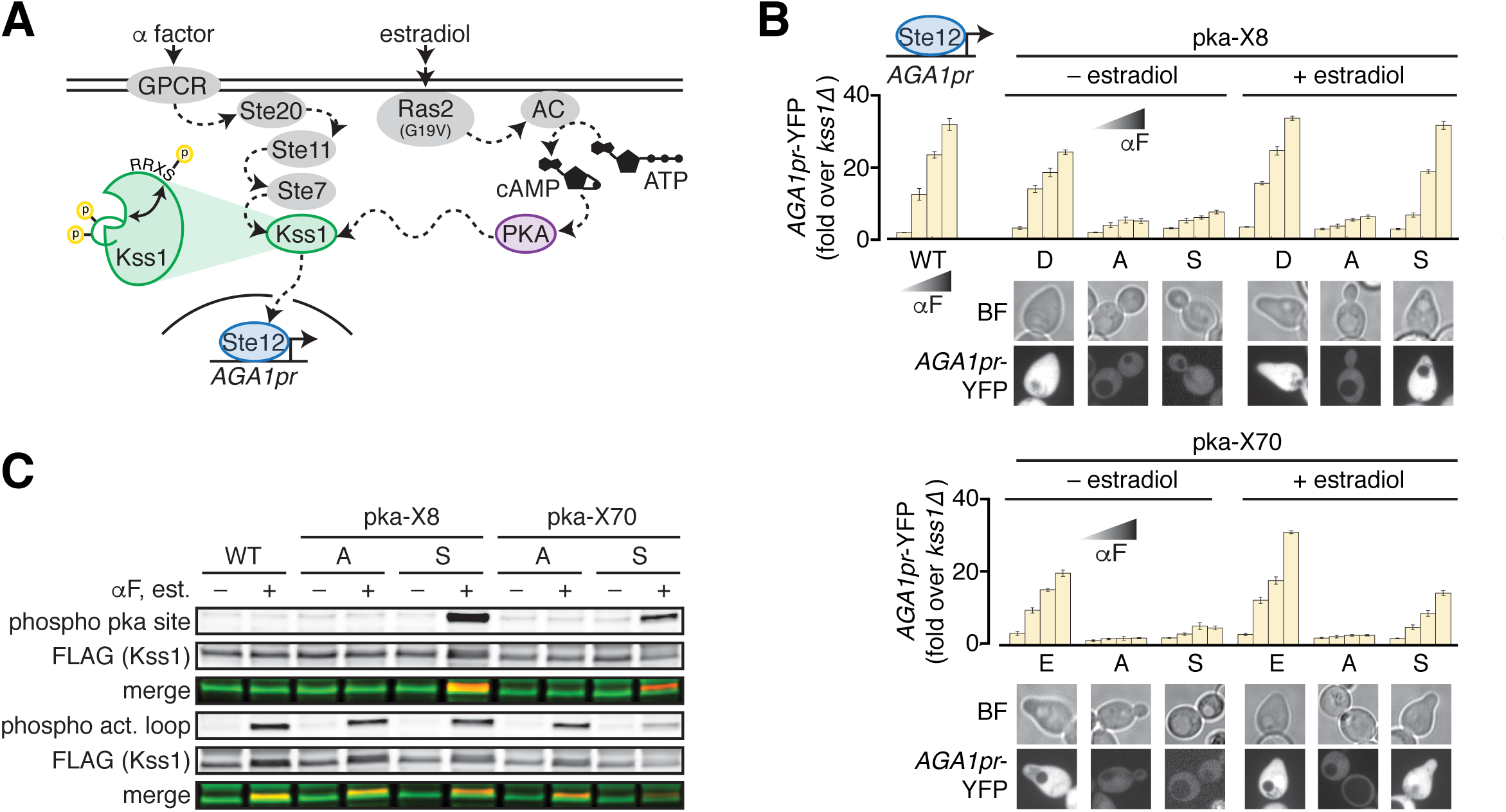
Engineering allosteric control of Kss1 by PKA phosphorylation. A.Cartoon of the engineered PKA‐ and Kss1-dependent yeast pheromone pathway. In this schematic, Kss1 activation requires both activation loop phosphorylation by the upstream MAP2K Ste7, and phosphorylation by PKA at an allosterically coupled surface. To experimentally increase PKA activity, expression of constitutively activated Ras2(G19V) is induced by addition of estradiol, which in turn activates adenylate cyclase (AC) to generate cyclic AMP (cAMP) from ATP to activate PKA. **B.** Kss1 mutants with PKA phosphorylation site consensus motifs introduced near position 8 (pka-**X**8, upper panel) or position 70 (pka-**X**70, lower panel) were assayed for expression of the *AGA1pr-YFP* reporter as in Fig. 2c. “**X**” stands for the amino acid at position 8 or 70 as denoted under the bar graphs. The images below the bar graphs show morphology and expression of the *AGA1pr-YFP* reporter in yeast cells bearing the indicated Kss1 mutants in the presence of 1 μM alpha factor following growth in the presence or absence of 20 nM estradiol. The data indicate that phosphorylation by PKA at these positions can allosterically regulate Kss1 activity.**C.** 3xFLAG-tagged wild type Kss1 and pka-**X**8 and ‒**X**70 mutants were immunoprecipitated from untreated cells or cells that had been treated with both 20 nM estradiol and 1 μM alpha factor. IP eluates were analyzed by Western blotting for total Kss1 as well as Kss1 phosphorylated on its activation loop (phospho act. loop) or at the engineered PKA site (phospho pka site). Merged images show that all mutants can be phosphorylated on their activation loop in the presence of alpha factor, but only pka-S8 and pka-S70 can be phosphorylated by PKA in the presence of estradiol.

Introduction of pka-E at position 68 resulted in Kss1 loss-of-function (Supplementary Fig. 1E, F), indicating that in this instance, the mutation of positions 65-66 to arginine to introduce the PKA site was not neutral. However, introducing the PKA consensus motif at positions D8 and E70 showed the complete expected pattern of activity for gain of phosphoregulation (Fig. 3B). For both sites, introduction of the two arginine residues upstream was near neutral, mutation of the negatively charged residue caused loss of function, and Kss1 pka-S activity depended on enhanced PKA activity via estradiol-induced expression of Ras2^G19V^ (Fig. 3B). Immunoprecipitation of the 3xFLAG-tagged Kss1 mutants followed by Western blot analysis supports this finding. Both the pka-A and pka-S variants displayed activation loop phosphorylation when treated with alpha factor, indicating that they remain substrates of Ste7.

However, only Kss1-pka-S8 and Kss1-pka-S70 were recognized by an antibody specific for phosphorylated PKA substrates when purified from cells treated with estradiol (Fig. 3C). Moreover, both Kss1-pka-S8 and Kss1-pka-S70 were able to induce the morphological response to pheromone – the mating projection known as the “shmoo” – in an αF‐ and PKA activity-dependent fashion (Fig. 3B). Thus, the transcriptional and physiological outputs of Kss1 can be rewired to depend on an orthogonal input by a stepwise process of introducing a phosphorylation site at latent allosteric surface sites.

What is special about the nine surface negatively charged amino acids that they are allosterically pre-wired to regulate the Kss1 active site? Is this functional coupling idiosyncratic to Kss1 or conserved in the kinase family? To address this, we used the Statistical Coupling Analysis (SCA) (*22-24*) to examine the correlated conservation (or coevolution) of amino acid positions in an alignment encompassing all EPK subfamilies and a focused alignment of the CMGC subfamily that includes the MAP kinases (Supplementary Fig. 2A,B). The basic result from SCA is the finding that protein families have internal networks of coevolving amino acids (called “sectors”) that tend to link protein active sites to distantly positioned allosteric surface sites (*22, 25-28*). Consistent with this, we identified a protein sector in the EPK family that forms a physically contiguous network of amino acids within the three-dimensional structure (Supplementary Fig. 2C-E, Supplementary Table 4). The sector is enriched for positions associated with kinase function: comparison to a deep mutational scan of human ERK2 (29) shows a clear, statistically significant association between the sector and sites associated with loss-of-function (*p* = 2.3E-19 by Fisher Exact Test, Supplementary Fig. 3, Supplementary Table 5). Further, the sector encapsulates several structural motifs well known to be associated with kinase activation including the αC-helix, the DFG motif and the catalytic and regulatory spines (Supplementary Fig. 2C-E, Supplementary Fig. 4) (*30, 31*). Thus, like for other proteins, analysis of conserved coevolution of amino acids in EPKs provides a sparse, distributed model for the functionally relevant energetic connectivity of amino acids (*24*).

**Figure 4.**
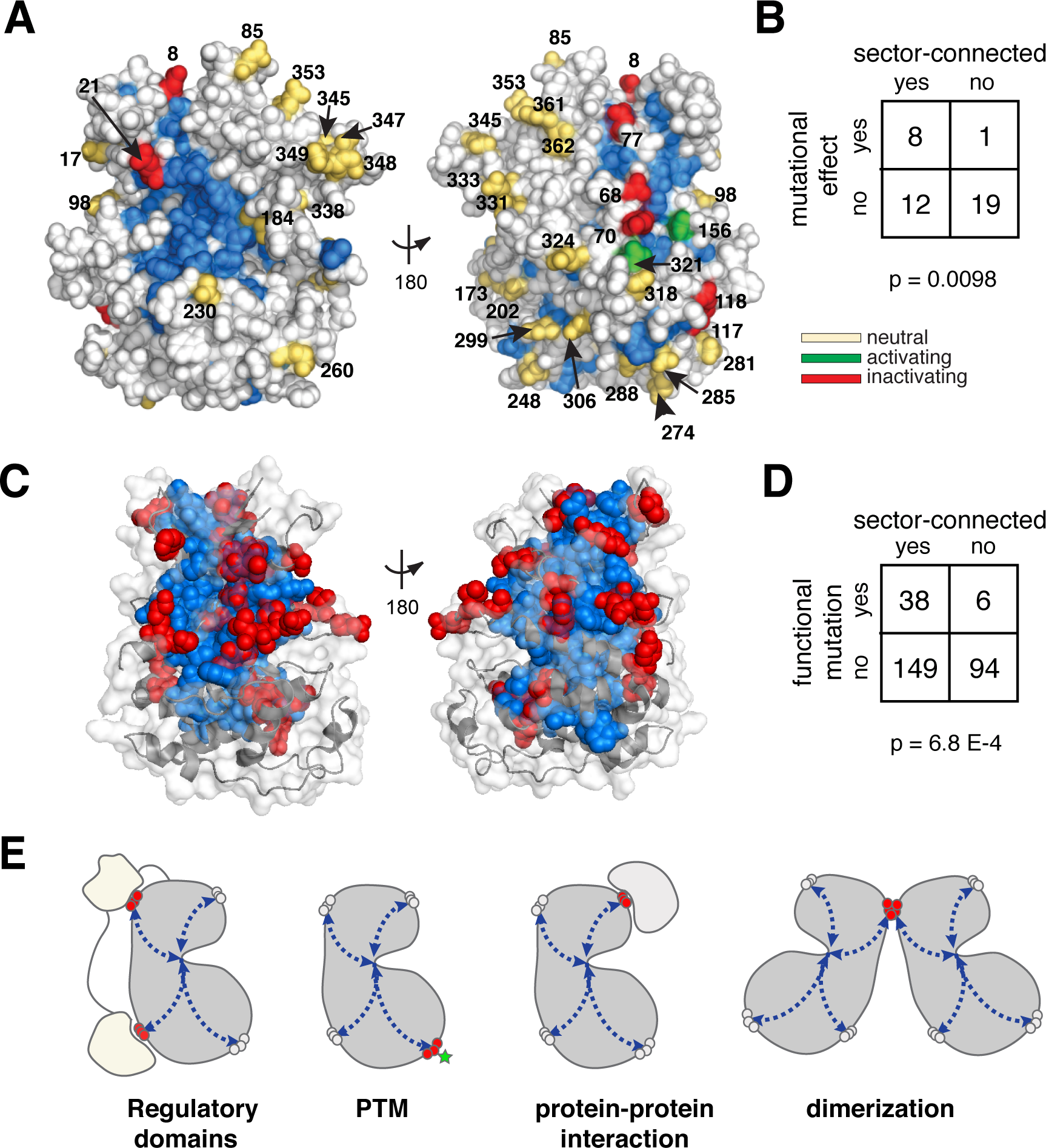
Sector connected surface sites are hotspots for allosteric regulation. **A.** Space filling diagram of a Kss1 homology model (*34*). The CMGC sector, defined as positions that co-evolve across the CMGC kinases, is indicated in blue. Acidic surface residues with a neutral, activating, or inactivating effect on kinase function upon mutation to alanine are shown as yellow, green or red spheres respectively. **B.** Fisher’s exact table demonstrating statistically significant enrichment of acidic surface residues with a functional effect upon mutation at sector-connected positions. To be sector connected, a position must have at least one atom within 4 Å of the sector. **C.** The EPK superfamily-wide sector (blue spheres) mapped to the CMGC yeast kinase Pho85 (PDB: 2PK9, grey cartoon and surface). Red positions are sites collected from the literature known to alter kinase function when mutated in a functional study or human disease context (Supplementary Table 7). **D.** Fisher’s exact table demonstrating statistically significant enrichment of the functional mutations shown in **c** at sector-connected positions. **E.** Model for the evolution of regulatory diversity. Latent allosteric sites distributed across the protein surface (red circles) are connected to the active site via a protein sector (blue arrows). These sites are poised for the acquisition of new regulation via evolutionary, disease, or engineering processes. In any particular family member, only a subset of sites may be used, and the regulatory mechanism need not be conserved across homologs.

Previous work has proposed that the sector represents the physical mechanism that underlies the pre-wiring of surface sites that serve as hotspots for the emergence of new allosteric regulation (*10, 32*). Consistent with this, eight of the nine functionally coupled surface D/E residues in Kss1 are sector connected (p = 0.0098, Fisher Exact Test) (Fig. 4A, B), including the two that yield new PKA-dependent phosphoregulation. Thus, the gain of new regulatory function in Kss1 occurs at sites that are not idiosyncratic, but that interact with an allosteric network that coevolves in the entire kinase family. This result is robust to details of alignment construction and statistical cutoffs for determining sector positions (Supplementary Fig. 5, Supplementary Table 6). These data support a model that new regulation preferentially emerges in proteins at surface sites that are evolutionarily prewired in protein families.

If so, all natural kinases should follow the principle that functionally sensitive and physiologically relevant allosteric sites, regardless of mechanism, should be found with statistical preference at sector-connected surfaces. The sector-connected surfaces would then provide an explanation for the diversity of regulatory sites observed in extant kinases (Fig. 1). To investigate this, we constructed a curated database of mutations sampled across a diversity of kinases (those listed in Fig 1A, Supplementary Table 7). These mutations were selected because they were experimentally demonstrated to disrupt kinase regulation and/or function, and, in many cases, are also associated with disease. An analysis of mutations sampled across the kinase superfamily reveals a clear pattern: functional mutations cluster around the sector edges with strong statistical preference (p = 0.00068, Fisher Exact Test) (Fig. 4c, d, Supplementary Table 8). Thus, we conclude that the natural architecture of the protein kinases does indeed facilitate the evolution of regulatory diversity.

The central idea supported by our results is that proteins contain a conserved cooperative mechanism that endows specific sites on the protein surface with a latent capacity for allostery. As a consequence of this intrinsic cooperative architecture, allosteric regulation may emerge in a variety of mechanistic forms at multiple, distinct locations in different family members (Fig. 4E). Our results demonstrate a general strategy for engineering new cell signaling pathways – *in vivo* phospho-regulation can in principle be introduced into any soluble protein by targeting negatively charged residues at sector-connected surfaces (3). Further, the sector provides a context for interpreting kinase mutations involved in disease (Fig. 4C), and suggests possible cryptic sites for the development of allosteric inhibitors (8). Overall, this model provides a path for understanding how complex regulatory systems evolve, and suggests that sector edges provide a substrate for generating variation in cellular signaling and communication.

## ACKNOWLEDGEMENTS

This collaboration was initiated at the 2013 q-bio conference held at St. Johns College, Santa Fe, NM. We would like to thank R. Ranganathan for discussion and comments on the manuscript. We are grateful to the Whitehead Institute FACS facility and the Keck Microscopy facility for technical assistance. This work was supported by an NIH Early Independence Award (DP5 OD017941-01 to D.P.), the Green Center for Systems Biology, and the Gordon and Betty Moore Foundation’s Data-Driven Discovery Initiative (Grant GBMF4557 to K.R.).

## AUTHOR CONTRIBUTIONS

Conceptualization, K.A.R, D.P., and O.R.; Methodology, K.A.R., D.P., O.R. and P.C.; Investigation, D.P., J.P.P, O.R., and K.A.R., Writing – Original Draft, D.P. and K.A.R; Writing – Reviewing & Editing, D.P., J.P.P, O.R., and K.A.R, Supervision, D.P. and K.A.R.

## SUPPLEMENTARY MATERIALS

### Supplementary Figures 1-6

**Figure 1.** Experimental approach to introduce a PKA phosphorylation site that controls MAPK Kss1 activity

**Figure 2.** Statistical Coupling Analysis (SCA) of the Eukaryotic Protein Kinases (EPKs)

**Figure 3.** ERK2 mutations within the kinase sector are enriched for loss-of-function

**Figure 4.** The kinase sector encompasses the catalytic and regulatory spines

**Figure 5.** The relationship of negatively charged surface positions to the kinome-wide EPK sector

### Supplementary Tables 1-8

**Table 1.** Plasmids

**Table 2.** Yeast strains

**Table 3.** Comparison of Kss1 point mutations from the literature with our data Table 4. List of sector positions for several representative kinases

**Table 5.** Statisticalassociationbetween thesector,conservationandERK2 mutational data

**Table 6.** Statisticalassociationbetween thesector,conservationandKSS1 D/E surface mutations

**Table 7.** Curated set of functional mutationsfor a diversity of kinasesand references

**Table 8.** Statisticalassociationbetween thesector,conservationandfunctional mutations

sampled across a diversity of kinases

### Methods

## SUPPLEMENTARY FIGURE LEGENDS

**Supplementary Figure 1.**
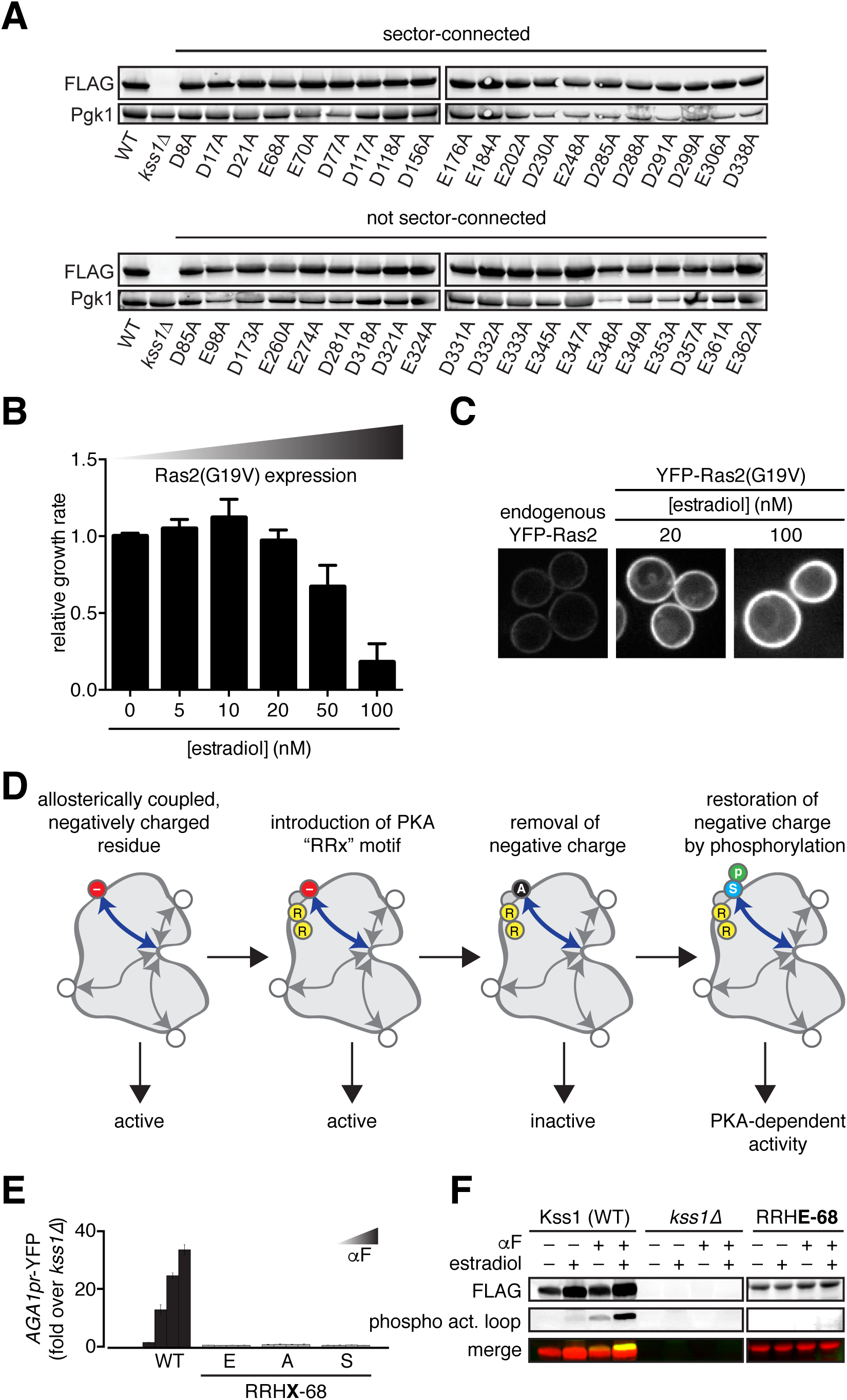
Experimental approach to introduce a PKA phosphorylation site that controls MAPK Kss1 activity. **A.** Expression levels for all forty Kss1 mutants in which each acidic surface position was mutated to alanine, alongside WT and *kss1*Δ cell controls. Kss1 (and all mutants) were tagged with 3xFLAG and expression level was monitored by anti-FLAG immunoblot. Total protein loaded in each lane was monitored with an anti-Pgk1 antibody. Mutations are grouped into sector-connected and not sector-connected categories, as discussed later in the text. All alanine mutants show expression levels similar to WT. **B.** To increase PKA activity, constitutively active Ras2(G19V) was expressed from a promoter activated in proportion to the concentration of estradiol in the media. Growth was monitored in log phase by measuring OD_600_ over time in cells treated with the indicated concentrations of estradiol and plotted relative to cells without estradiol. Error bars are the standard deviation of three independent cultures. Based on these data, we chose to use 20 nM estradiol for all experiments because this is the highest concentration that did not result in growth inhibition.**C.** Localization of YFP-Ras2 expressed from its endogenous promoter and YFP-Ras2(G19V) expressed at two concentrations of estradiol, showing significant over-expression at 20 nM but proper plasma membrane localization.**D.** Schematic of the mutational strategy to introduce a functional PKA phosphorylation site at an allosterically coupled negatively charged surface position (red circle with “-” sign). First, two consecutive Arg residues are introduced by mutation (RRx) at the ‐2 and ‐3 positions with respect to the negatively charged (Asp/Glu) position to create the PKA consensus motif. While the Asp/Glu is maintained at position 0, the kinase must retain function in the presence of the RRx. Next position 0 is mutated to Ala in the context of the RRx to remove the negative charge. Removal of the negative charge should result in a loss-of-function kinase. Finally, mutation of position 0 to Ser must conditionally restore kinase activity in the presence of PKA activity.**E.** Insertion of the RRx motif at position 68 resulted in Kss1 loss-of-function as assayed for activation of the *AGA1pr*-YFP reporter by flow cytometry following treatment with 0, 0.01, 0.1 and 1 μM αF for 4 hours. Bars represent the average of the median YFP fluorescence from 3 biological replicates normalized to the untreated *kss1*Δ cells, and error bars are the standard deviation of the biological replicates. **F.** Introduction of the RRx motif at position 68 resulted in reduced Kss1 protein expression and loss of activation loop phosphorylation. Samples of 3xFLAG-tagged WT and pka-E68 were monitored by anti-FLAG Western blot, and were also probed for phosphorylation of the activation loop (phospho act. loop)

**Supplementary Figure 2.**
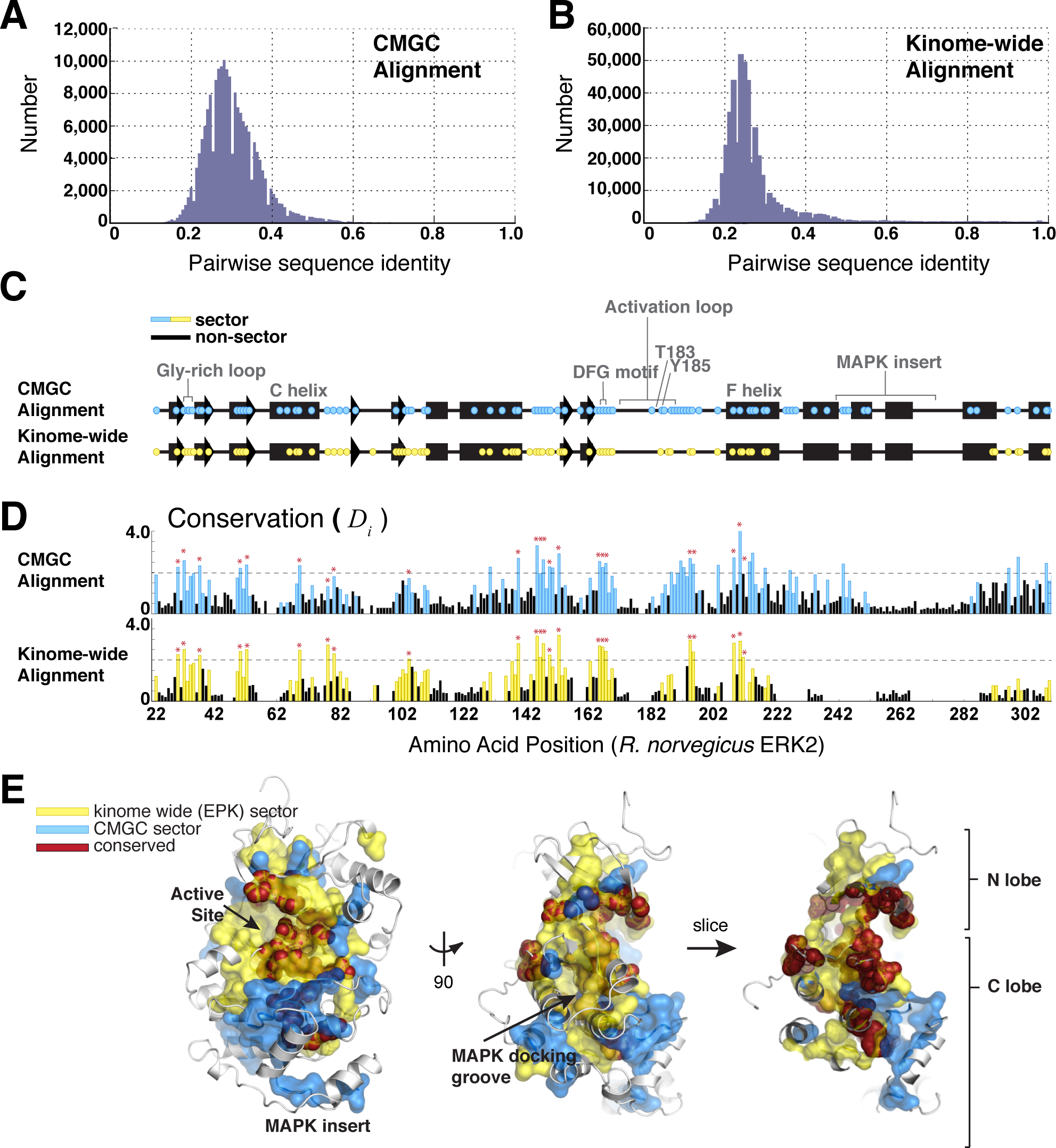
Statistical Coupling Analysis (SCA) of the Eukaryotic Protein Kinases. The analysis was performed for two different multiple sequence alignments of the kinase catalytic domain: one specific to the CMGC kinases (635 sequences), and one containing 7128 kinases sampled across the kinome. **A.** Histogram showing the distribution of pairwise sequence identities computed across all pairs of sequences in the CMGC alignment. **B.** As in (**A**) but for the kinome wide alignment. Both alignments show a unimodal distribution with a mean pairwise sequence identity near ~25%. **C.** Sector positions derived from the CMGC alignment (blue) or kinome-wide alignment (yellow) are distributed along the primary and secondary structure of the CMGC/MAPK ERK2. Subfamily-specific regions, such as the MAPK-insert, are only part of the sector derived from the CMGC alignment. **D.** The relationship between the sector and positional conservation (computed as the Kullback-Leibler relative entropy, *D*_*i*_) for both the CMGC and kinome-wide alignments. Sector positions are highlighted in blue or yellow for the CMGC and kinome-wide alignments respectively. Red stars indicate highly conserved positions (defined as *D*_*i*_ > 2.0 in the kinome-wide alignment). **E.** The kinome-wide and CMGC-specific sectors (yellow and blue transparent surfaces, respectively) mapped on human ERK2 (gray ribbon) (PDB: 2ERK). Conserved positions are shown as red spheres.

**Supplementary Figure 3.**
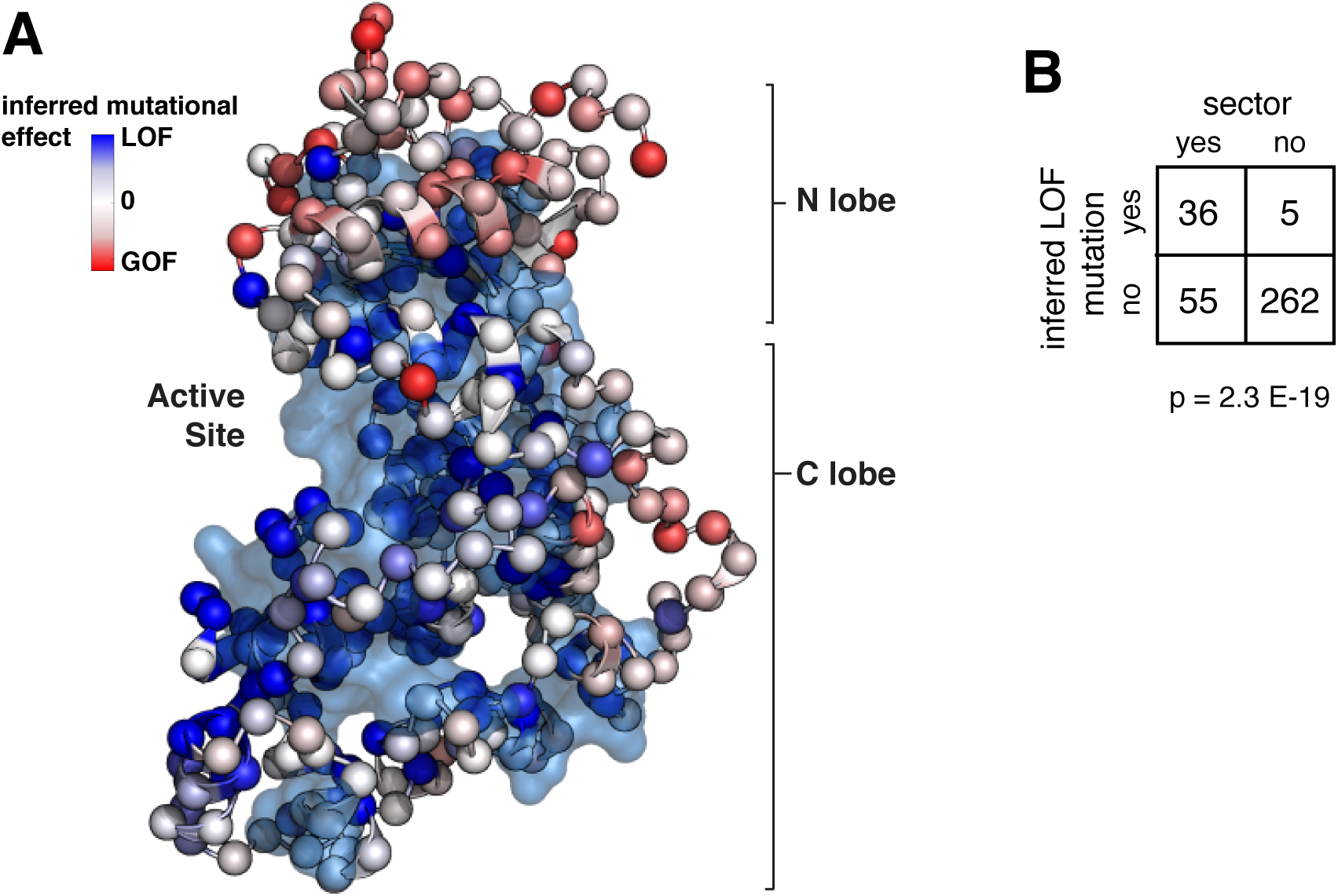
ERK2 mutations within the kinase sector are enriched for loss-of-function. **A.** The CMGC sector displayed as a transparent cyan surface overlaid by a ball-and-stick model of *R. norvegicus* ERK2 (PDB: 2ERK). Red-white-blue heat map color coding of the ball-and-stick model indicates residues that when mutated by Brenan et al. (29) led to inferred gain-of-function (GOF), neutral, and loss-of-function (LOF) activity, respectively, of human ERK2. **B.** Fisher’s exact table demonstrating statistically significant enrichment of inferred LOF mutations in ERK2 with CMGC sector-connected positions.

**Supplementary Figure 4.**
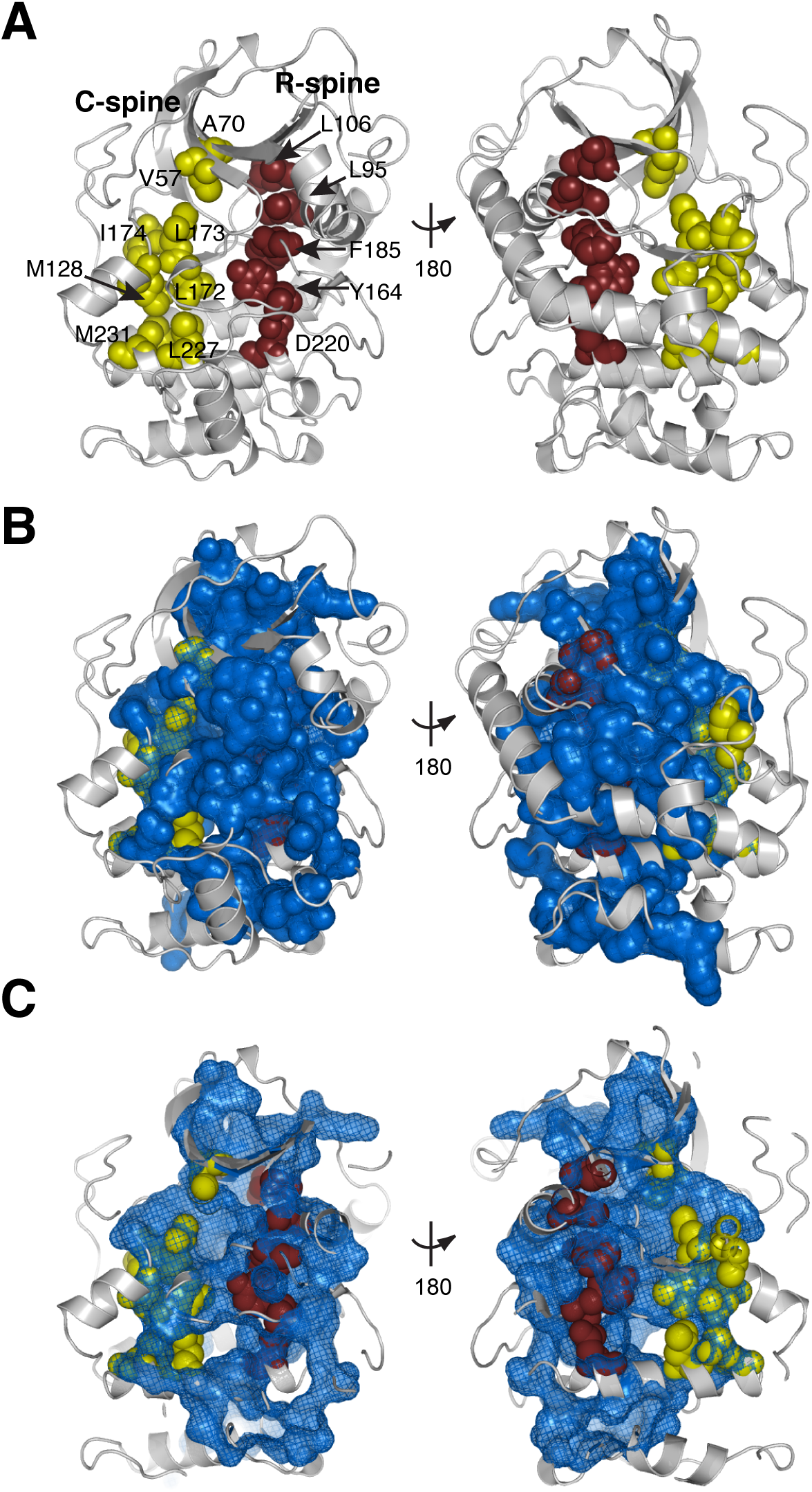
The kinase sector encompasses the catalytic and regulatory spines. **A.** The positions of the catalytic (C) and regulatory (R) spines as defined by Kornev et al (*30, 31*), yellow and dark red spheres respectively) are shown on a grey ribbon diagram of protein kinase A (PDB: 2CPK). **B.** The kinome-wide sector is overlaid in blue on panel (**A**). **C.**A vertical slice half way through panel (**B**) revealing the overlap between the spines and the sector. All spine positions are encapsulated within the sector or sector-connected.

**Supplementary Figure 5.**
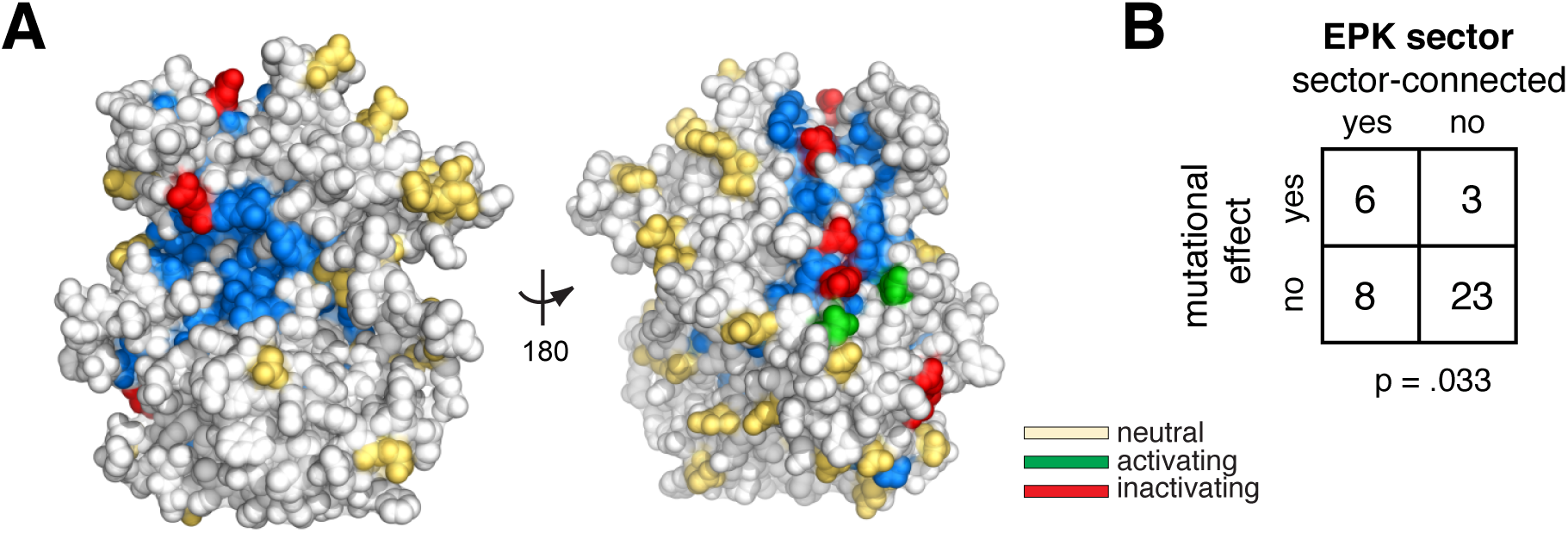
The relationship of negatively charged surface positions to the kinome-wide EPK sector. Within the main text, we show a statistically significant association with the sector defined using the CMGC specific alignment. For comparison, here we show the results with the kinome-wide (EPK) alignment. **A**. Space-filling model of Kss1 in gray, with the kinome-wide sector in blue. Positions with neutral, loss-of-function and gain-of-function mutations are color-coded (yellow, red, green, respectively). **B**.Fisher’s exact table demonstrating statistically significant enrichment of functional residues at sector-connected positions derived from the kinome-wide alignment.

## METHODS

### Yeast strains and plasmids

Yeast strains and plasmids used in this work are described in Supplementary Tables 1 and 2, respectively. All strains are in the W303 genetic background. Gene deletions were performed by one-step PCR as described (*35*). All Kss1 mutants were integrated into yeast genome as a single copy expressed from the endogenous *KSS1* promoter.

### Site-directed mutagenesis

Site-directed mutagenesis was performed with QuickChange according to the manufacturer’s directions (Agilent).

### Cell growth and treatment with α factor

All cells were grown in synthetic complete media with dextrose (SDC). Three single colonies from each Kss1 strain bearing the *AGA1pr-YFP* reporter were inoculated in 1 ml SDC in 2 ml 96-well deep well plates and serially diluted 1:5 three times. Plates were incubated overnight at 30°C. In the morning cells from the row that had been diluted 1:25 were typically found to have OD_600_ ~0.5. These cells were diluted 1:5 in 4 rows of a 96 well U-bottom micro-titer plate in a total volume of 180 μl and incubated for 1 hour at 30°C. In each row, cells were treated with different concentrations of α factor: 0, 0.01, 0.1 and 1 μM (10× stocks of α factor were prepared and 20 μl were added to 180 μl cells). Treated cells were incubated for an additional 4 hours at 30°C before translation was stopped by addition of 50 μg/ml cycloheximide. Cells were incubated for an additional hour at 30°C to allow time for fluorophores to mature. For experiments with estradiol, everything is the same except that all media contained 20 nM estradiol for the duration of the overnight growth and throughout the experiment.

### Flow cytometry

The AGA1pr-YFP reporter was measured by flow cytometry by sampling 10 μl of each sample using a BD LSRFortessa equipped with a 96-well plate high-throughput sampler. Data were left ungated and FlowJo was used to calculate median YFP fluorescence. Bar graphs show the average of the median of the three independent colonies that were assayed, and error bars are the standard deviation.

### Confocal microscopy

96 well glass bottom plates were coated with 100 μg/ml concanavalin A in water for 1 hour, washed three times with water and dried at room temperature. 80 μl of cells that had been treated with pheromone at the indicated concentrations for 3 hours were diluted to OD_600_ ~0.05 and added to a coated well. Cells were allowed to settle and attach for 15 minutes, and unattached cells were removed and replaced with 80 μl SDC media. Imaging was performed at the W.M Keck Microscopy Facility at the Whitehead Institute using a Nikon Ti microscope equipped with a 100×, 1.49 NA objective lens, an Andor Revolution spinning disc confocal setup and an Andor EMCCD camera. Images were analyzed in ImageJ.

### Immunoprecipitation of 3xFLAG-tagged Kss1 and mutants

2× 250 ml cultures of each strain were grown to OD_600_=0.8 at 30°C with shaking, one in SDC and the other in SDC + 20 nM estradiol. The SDC culture was left untreated while the SDC + estradiol culture was treated with 1 μM alpha factor for 30 minutes. Samples were collected by filtration and filters were snap frozen in liquid N_2_ and stored at ‐80°C. Cells were lysed frozen on the filters in a coffee grinder with dry ice. After the dry ice was evaporated, lysate was resuspended in 1 ml IP buffer (50 mM Hepes pH 7.5, 140 mM NaCl, 1 mM EDTA, 1% triton ×-100, 0.1% DOC, complete protease inhibitors), transferred to a 1.5 ml tube and spun to remove cell debris. Clarified lysate was transferred to a fresh tube and serial IP was performed. First, 25 μl of anti-FLAG magnetic beads (50% slurry, Sigma) were added, and the mixture was incubated for 2 hours at 4°C on a rotator. Beads were separated with a magnet and the supernatant was removed. Beads were washed 5 times with 1 ml IP buffer and bound material eluted 2x with 25 μl of 1 mg/ml 3xFLAG peptide (Sigma) in IP buffer by incubating at room temperature for 10 minutes. Beads were separated with a magnet and the two eluates were pooled in a fresh tube. 10 μl eluate was analyzed by Western blotting.

### Western blotting

Total protein was TCA purified from cells as described (6). 10 μl of each sample was loaded into 4-15% gradient SDS-PAGE gels (Bio-Rad). Gels were run at 25 mA for 45 minutes, and blotted to PVDF membrane at 225 mA for 40 minutes. After 1 hr blocking in Li-Cor blocking buffer, membranes were incubated with anti-FLAG primary antibody (SIGMA, F3165), anti‐phospho-PKA substrate, anti-phospho p44/42 (Cell Signaling, 9101), and/or anti-PGK (22C5D8) overnight at 4°C on a platform rotator (all 1:1000 dilutions in blocking buffer). Membranes were washed three times with TBST and probed by anti-mouse or anti-rabbit IR dye-congugated IgG (Li-Cor, 926-32352, 1:10000 dilution). The fluorescent signal was detected with the Li-Cor/Odyssey system.

### Statistical Coupling Analysis (SCA)

SCA was performed as described in (24) using PySCA 6 (http://reynoldsk.github.io/pySCA/) for two different multiple sequence alignments of the kinase catalytic domain: one specific to the CMGC kinases (635 sequences), and one containing 7128 kinases sampled across the kinome. The CMGC alignment was constructed by searching kinbase (http://kinase.com/kinbase/). Sequences were filtered for a length of 250-350 amino acids, and aligned by Promals3D (37) including the PDBS: 2B9H, 1BI8, 1Q97, 2ERK, 2F49, 2F9G, 2IW8, 2R7I, as reference structures. The kinome-wide alignment was previously constructed by the Shokat lab and was downloaded from http://sequoia.ucsf.edu/ksd/ (38). Following alignment processing and the application of sequence weights (as described in (*24*)), the alignments contained 464 and 380 total effective sequences for the CMGC and EPK alignments respectively. For both alignments, we followed an identical procedure for defining the sector. Briefly, we compute a conservation-weighted covariance matrix between all pairs of amino acid positions (see Supplementary Text for discussion of the relationship between the sector, conservation, and allosteric hotpots). This matrix provides a statistical description of the "evolutionary coupling" between all pairs of amino acid positions. We then analyze this matrix by conducting principle components analysis (PCA), and rotating the top eigenmodes using independent components analysis (ICA). The top independent components are used to define sectors. For both kinase alignments, we define a single sector that includes all positions contributing to the top 4 independent components (ICs). The group of positions contributing to each IC groups is defined by fitting an empirical statistical distribution to the ICs and choosing positions above a defined cutoff (default, > 95% of the CDF). The full analysis of both families can be downloaded from github.

### Defining sector-connected solvent accessible surface sites

We computed the relative solvent accessible surface area (RSA) over a homology model of Kss1 (34) using Michel Sanner's MSMS with a probe size of 1.4 Å, excluding all water and heteroatoms (39). A cutoff of 20% RSA was used to define solvent exposed surface positions (10). "Sector-connected" is defined as a position where any atom is within 4.0Å of a sector position.

## SUPPLEMENTARY TEXT

### The relationship between the sector, conservation, and allosteric hotpots

In this work we analyze the statistical association between the sector and functional measurements of mutational effects from three different datasets: (1) saturation mutagenesis of ERK2 (6,810 mutations across 359 positions, Supplementary Fig. 3 and Supplementary Table 5) (29), (2) an alanine scan of negatively charged positions on the Kss1 surface (40 mutations, Fig. 4a,b, Supplementary Fig. 5, Supplementary Table 6) and (3) functional mutations across a diversity of kinases surveyed from the literature (78 mutations mapped to 45 unique sites, Fig. 4c,d, Supplementary Table 7-8). The goal of this section is to provide a more complete discussion of the sector definition, as well as the relationship between the sector and these experimental datasets.

The sector is defined using the top four independent components (ICs) of the so-called “SCA matrix” (*C̃*_*ij*_), a conservation-weighted covariance matrix between all pairs of amino acid positions (22, 24). In our analysis of the protein kinases, we group all of the positions contributing to the top ICs into a single sector, with the rationale that this is the most conservative interpretation in the absence of experimental data indicating functional or structural independence between residue groups. Though distinct from the goals of this paper, a further analysis of how parts of the sector may diverge in particular kinase subfamilies and how these residue groups relate to the evolutionary tuning of different biochemical properties is interesting, and addressed separately in concurrent work from Creixell et al (manuscript in prep.).

The conservation weighting of amino acid correlations is a defining feature of SCA and is applied with two complementary goals in mind: (1) to emphasize co-evolution between conserved (and thus likely functionally relevant) positions and (2) to minimize the contribution of purely phylogenetic correlations that are expected to emerge at weakly conserved positions.

The origin of the conservation weights is described more completely in (24), but the weights are applied as: 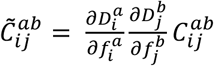, where *D*_*i*_ is the Kullback-Leibler relative entropy, and 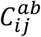 is the unweighted covariance between the frequencies (*f*) of a pair of amino acids *a*,*b* at a particular pair of positions *i*,*j*. Thus, we expect some redundancy between the information captured by sector positions (defined from the conservation weighted pairwise correlations) and simpler measures like the single-site conservation (*D*_*i*_) (*40*). Accordingly, we consider the statistical association of conservation alone with the functional mutagenesis data alongside our analysis of sector positions.

We computed the statistical association between functional mutations and either: (1) the sector, at several cutoffs (p = 0.95, 0.96, 0.97 and 0.98) and (2) conserved positions, at several cutoffs (*D*_*i*_ = 1, 1.15, 1.3 and 2). This was done for two alignments: one containing only the CMGC family kinases, and one encompassing kinases across the full kinome (referred to as the EPK alignment). The cutoffs for the sector were chosen to span a range of sector sizes (e.g. from 41-91 amino acid positions, ~15-33% of the kinase). The cutoffs for conservation (*D*_*i*_) were chosen to give similar numbers of amino acid positions as the sector cutoffs (Supplementary Table 5). These cutoffs include amino acid positions spanning “moderate” to more “stringent” levels of conservation: to map *D*_*i*_ to a more easily interpreted measure we computed the frequency of the most conserved amino acid at each position included in the cutoff. For the EPK alignment, we see that the *D*_*i*_ cutoffs of 1,1.15, 1.3 and 2 correspond to conserved amino acid frequencies of 0.23, 0.39, 0.39 and 0.68 respectively. Following the definition of sector and conserved positions, we used a one-tailed Fisher exact test on a two-by-two contingency table (as in Fig. 4b) to evaluate the probability that the observed association between the sector and experimental data (or any association more extreme) is obtained randomly. We found that both the sector positions and conserved positions have a statistically significant association with the functional data over a range of cutoffs (*p* < 0.05, Supplementary Tables 5,6,8). This observation forms the core of the argument that specific, evolutionarily conserved and co-evolving positions act as allosteric hotspots on the protein surface.

Though this statistical association does not depend strongly on the choice of alignment, we do observe some subtle differences. For example, the CMGC-specific alignment has a slightly better association with the ERK2 saturation mutagenesis data than the full alignment (Pearson χ^2^ p= 0.05), while the full alignment performs better than the CMGC alignment when compared to the kinome-wide sampling of mutations (Pearson χ^2^ *p* =4.7E-7). In both cases, the sector definition agrees better with the experimental data when the underlying alignment is more representative of the kinases being compared.

The sector positions and conservation show a statistically equivalent association with the functional data (as assessed by comparing the two contingency tables by Pearson χ^2^), meaning that it is difficult to distinguish between the functional significance of conserved residues and sector residues (40). However, the goal of this work is not to test the sector as an exclusive model for allosteric networks in proteins. Rather, our central claim is that allosteric potential is non-uniformly loaded into a handful of positions on the protein surface and that these facilitate the evolution of new regulation. The sector provides one way to identify these positions, and unlike single-site conservation, leads naturally to the interpretation that these positions form a cooperative network embedded within the protein structure.

### Background on Kss1

The MAPK Kss1 is expressed in both haploid and diploid *S. cerevisiae* cells. Kss1 is activated via phosphorylation by Ste7 (MEK). When overproduced, Kss1 stimulates recovery from pheromone-imposed G1 arrest and was first identified as a suppressor of Sst2 mutations (*16*). Kss1 is also involved in filamentous (invasive) growth in haploid cells (17) and pseudohyphal development in diploid cells (20). While Kss1 is concentrated in the nucleus, stimulation with mating pheromone results in relocation of Kss1 to the cytoplasm (*41, 42*).

▪ Length: 368 amino acids
▪ Kinase domain: residues 13-313
▪ ATP binding signature: residues 19-43
▪ Kss1 forms an initial tight complex with the MEK Ste7 (K_D_ of ~5 nM) that is not the ES conformation (43).
▪ Residues on Kss1 phosphorylated by the MEK Ste7 within the activation loop: T183, Y185
▪ The mutant K42R inactivates Kss1 activity, but does not affect phosphorylation of the activation loop residues.
▪ Kss1 binds to Ste7 (MEK), Ste12 (transcription factor), Dig 1 (transcription regulator, Kss1 substrate), Dig2 (transcription regulator, Kss1 substrate) and other phosphorylation substrates.
▪ Kss1 exhibits both a kinase-dependent positive activity and a kinase-independent inhibitory activity:
  ∘ Kss1 positive activity requires both activation by Ste7 and Kss1 catalytic activity,
  ∘ Kss1 inhibitory activity requires only the Kss1 protein

---

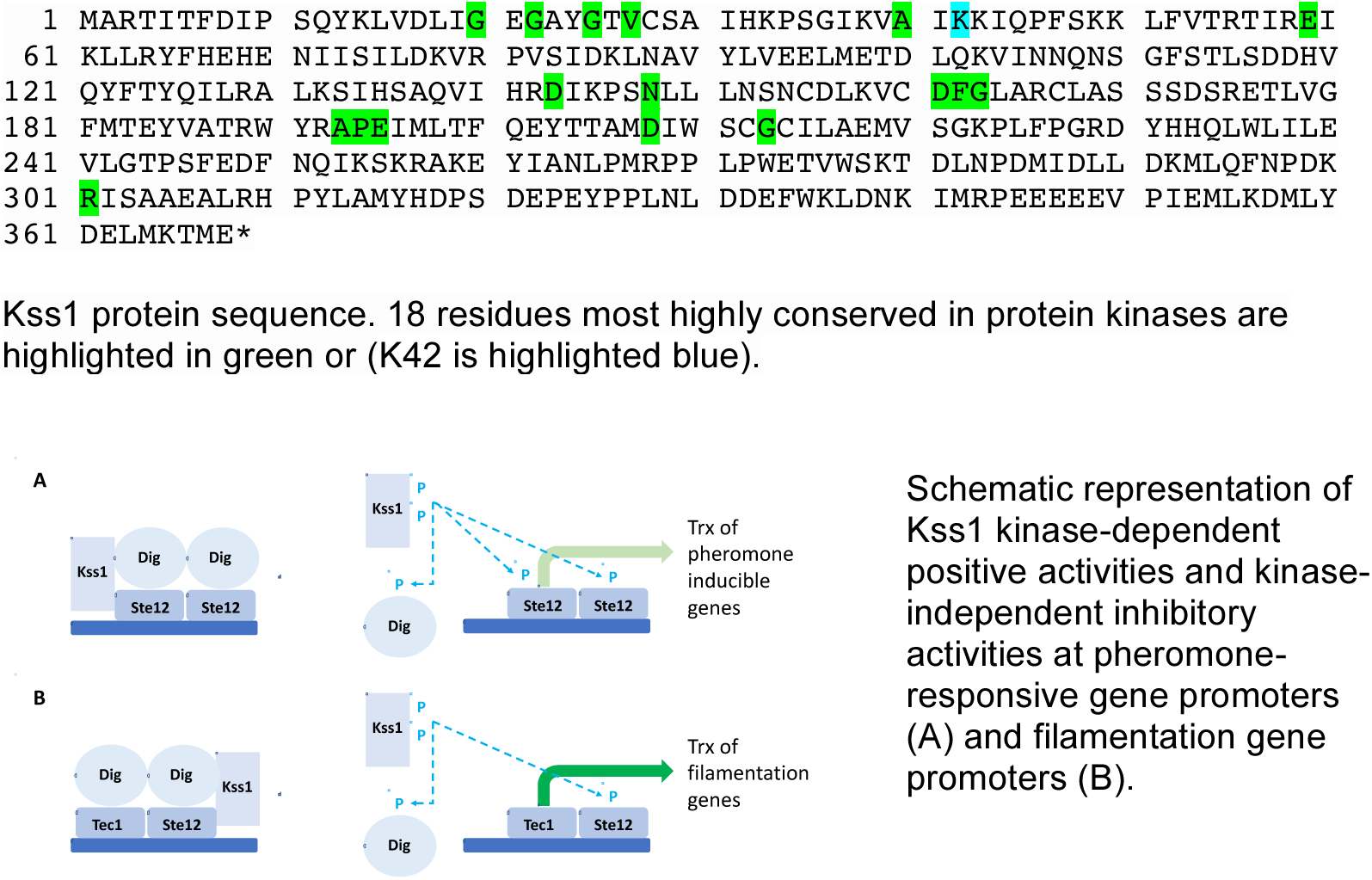

**Supplementary Table 1.** Plasmids

| pDP number | Nickname | Description | Marker |
| --- | --- | --- | --- |
| 11 | AGA1pr-YFP | pNH605-AGA1pr-YFP; single integration | *LEU2* |
| 436 | Kss1-3xFLAG-V5 | pNH604-KSS1pr-KSS1-3xFLAG-V5; single integration; template for all Kss1 mutants | *TRP1* |
| 1 | GEM | pRS306-GAL4dbd-EstradiolReceptor-VP16ad; integrative | *URA3* |
| 603 | Ras2(G19V) | pNH603-GALpr-RAS2(G19V) | *HIS3* |
| 333 | YFP-Ras2(G19V) | pNH603-GALpr-YFP-RAS2(G19V) | *HIS3* |

**Supplementary Table 2.**
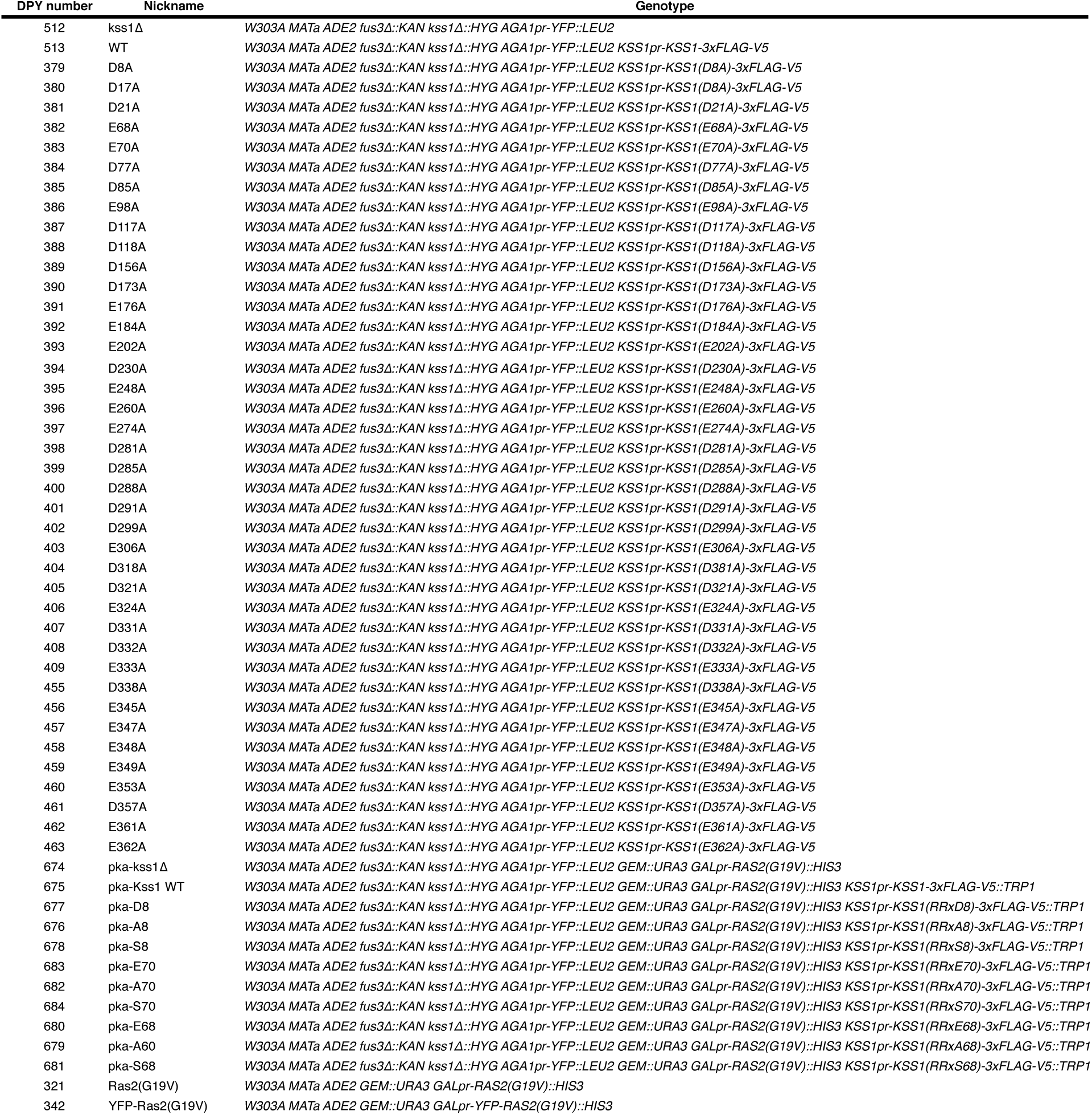
Yeast strains

**Supplementary Table 3.**
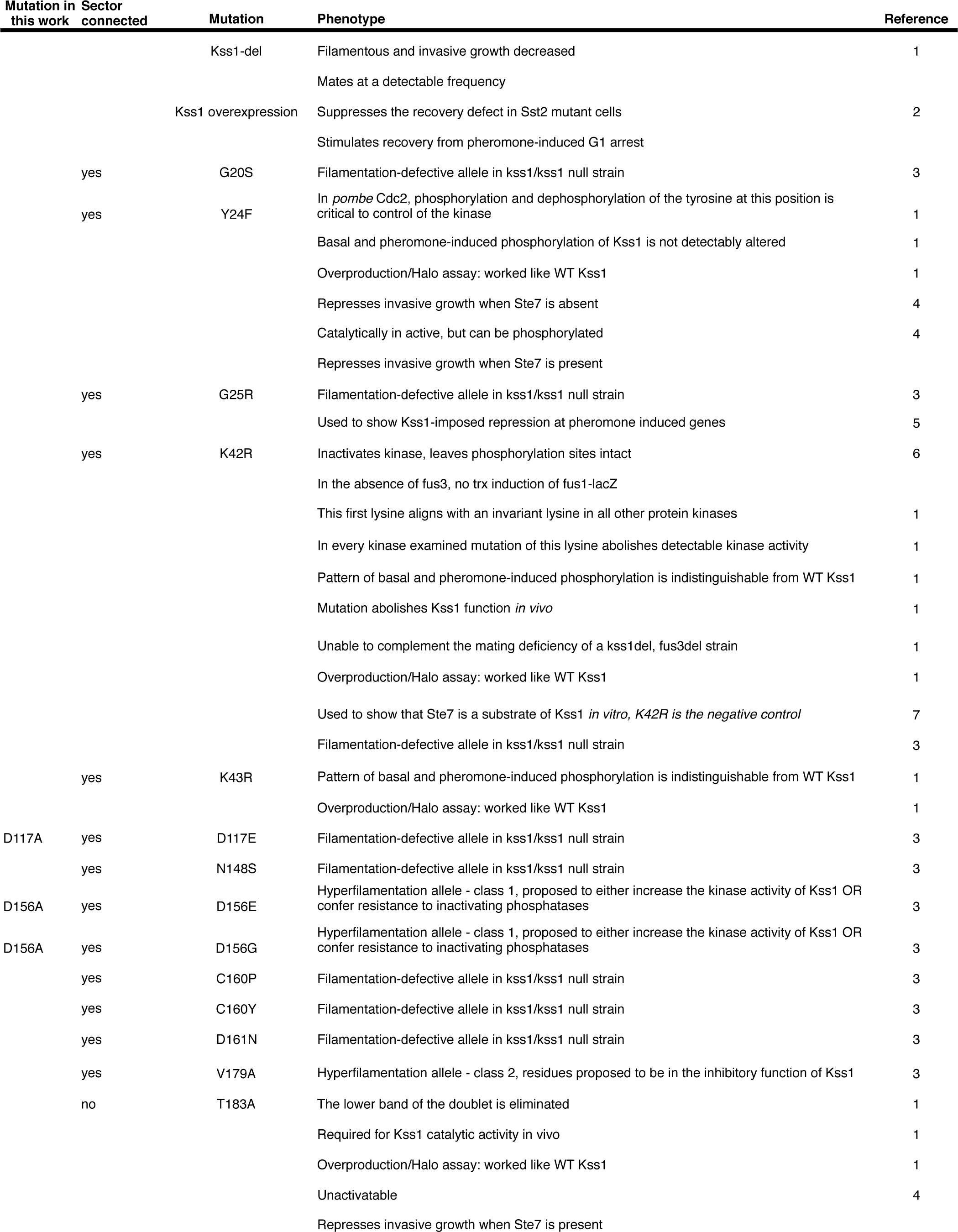
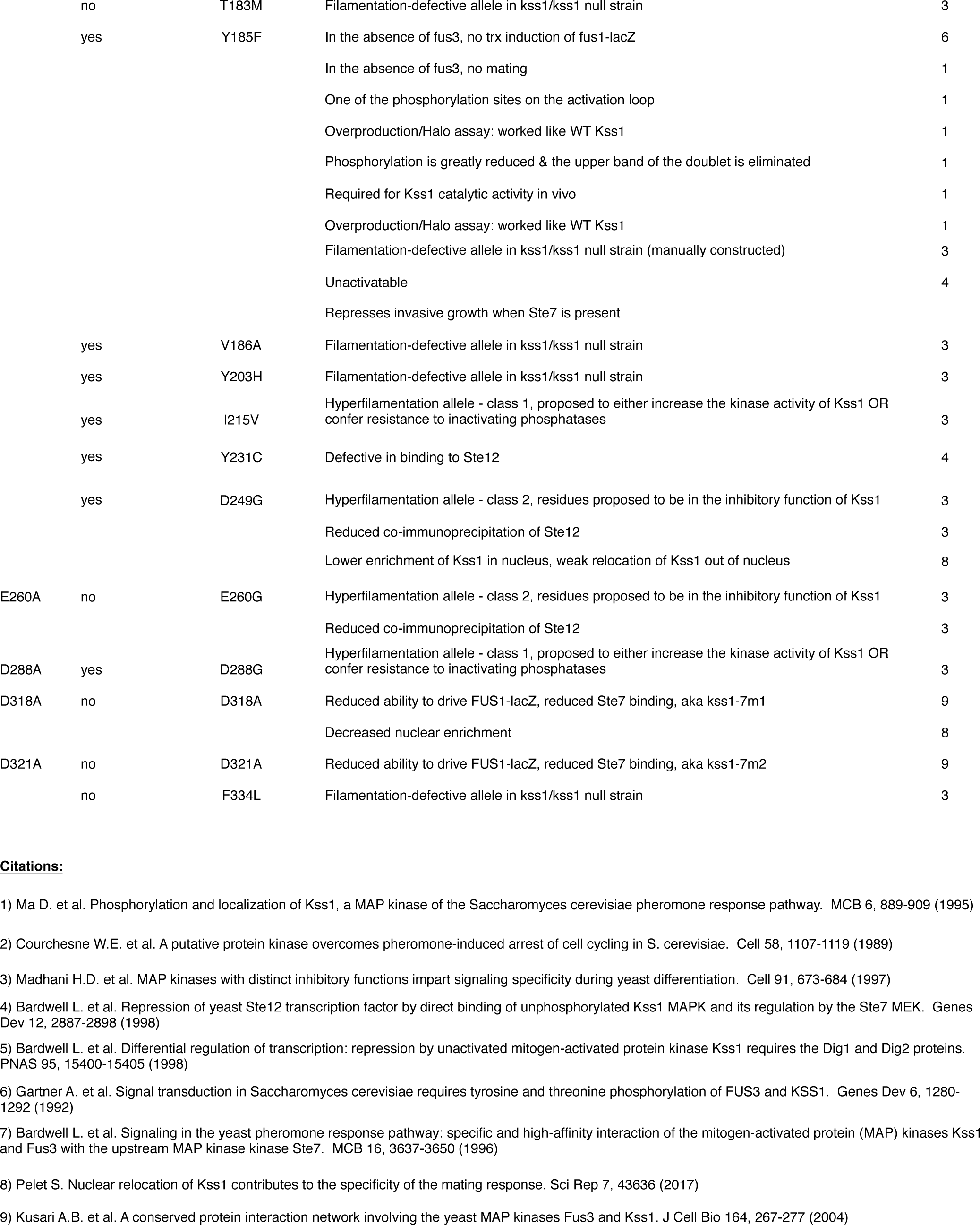
Comparison of Kss1 point mutations from the literature with our data and the CMGC sector (cutoff p=0.95).

**Supplementary Table 4.**
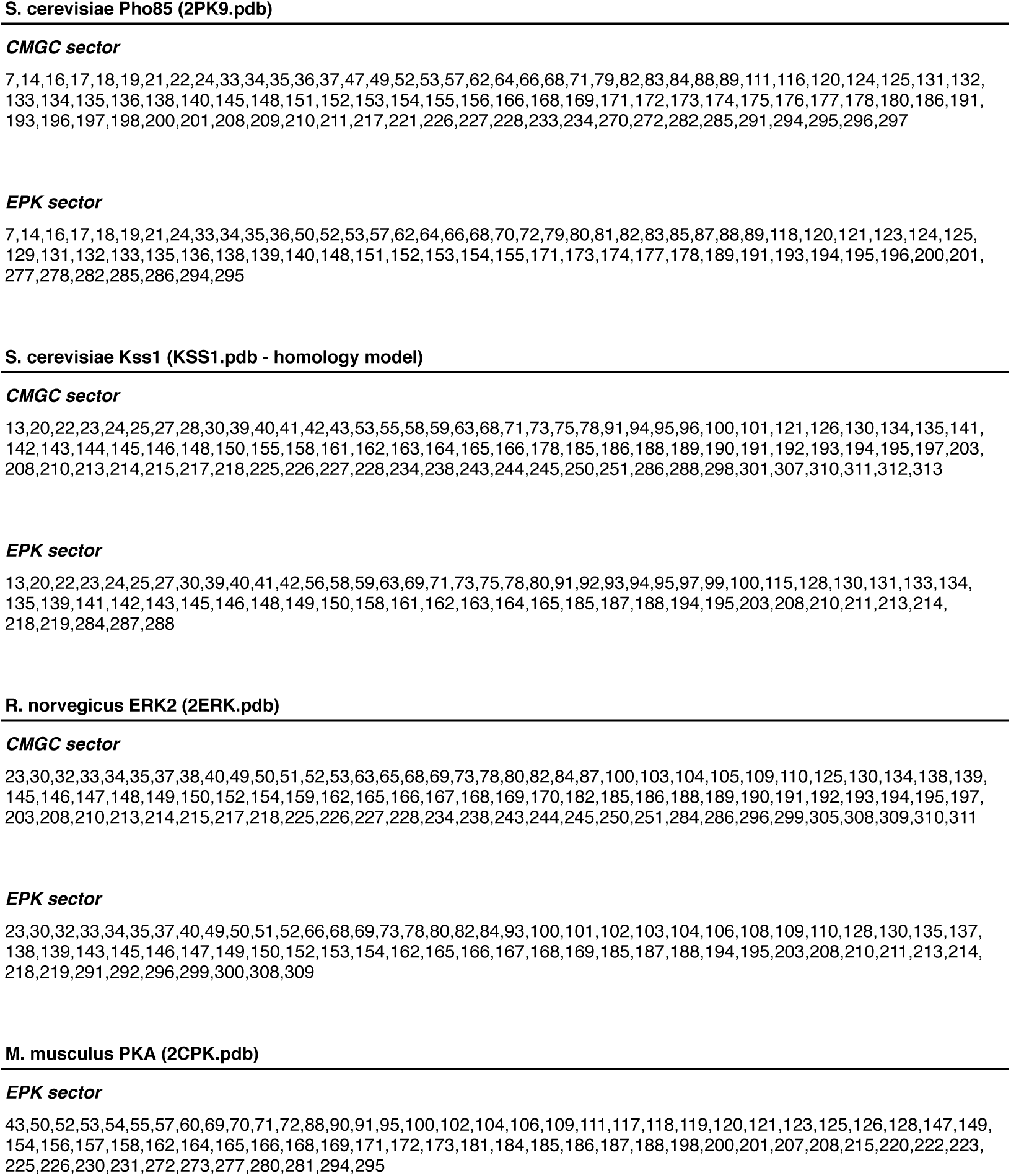
Sector positions mapped to several representative kinase structures

**Supplementary Table 5.** Statistical association between the sector, conservation and ERK2 mutational data

**Sector positions**
| sector cutoff: | 0.95 | 0.96 | 0.97 | 0.98 |
| --- | --- | --- | --- | --- |
| **CMGC alignment** | 2.3 E-19 (Npos = 91) | 4.4 E-20 (Npos = 81) | 1.4 E-20 (Npos = 62) | 8.4 E-10 (Npos = 41) |
| **EPK alignment** | 5.3 E-11 (Npos = 72) | 5.7 E-11 (Npos = 61) | 9.0 E-11 (Npos = 47) | 2.8 E-4 (Npos = 19) |

**Conserved positions**
| Di cutoff: | 1 | 1.15 | 1.3 | 2 |
| --- | --- | --- | --- | --- |
| **CMGC alignment** | 2.8 E-16 (Npos = 116) | 3.9 E-17 (Npos = 88) | 2.4 E-18 (Npos = 76) | 3.4 E-18 (Npos = 32) |
| **EPK alignment** | 4.7 E-8 (Npos = 72) | 4.2 E-8 (Npos = 60) | 3.7 E-10 (Npos = 44) | 1.5 E-10 (Npos = 23) |

**Supplementary Table 6.** Statistical association between the sector, conservation and KSS1 D/E Surface Mutations (*italics indicates insignificant relationships*)

**Sector positions**
| sector cutoff: | 0.95 | 0.96 | 0.97 | 0.98 |
| --- | --- | --- | --- | --- |
| CMGC alignment | 9.83E-03 | 6.14E-03 | *7.20E-02* | *1.38E-01* |
| EPK alignment | 3.29E-02 | 2.07E-02 | *1.90E-01* | *8.97E-01* |

**Conserved positions**
| Di cutoff: | 1 | 1.15 | 1.3 | 2 |
| --- | --- | --- | --- | --- |
| CMGC alignment | 1.52E-02 | 2.01E-02 | 2.01E-02 | *5.04E-01* |
| EPK alignment | *1.03E-01* | 3.28E-03 | 3.28E-03 | 8.50E-03 |

**Supplementary Table 7.**
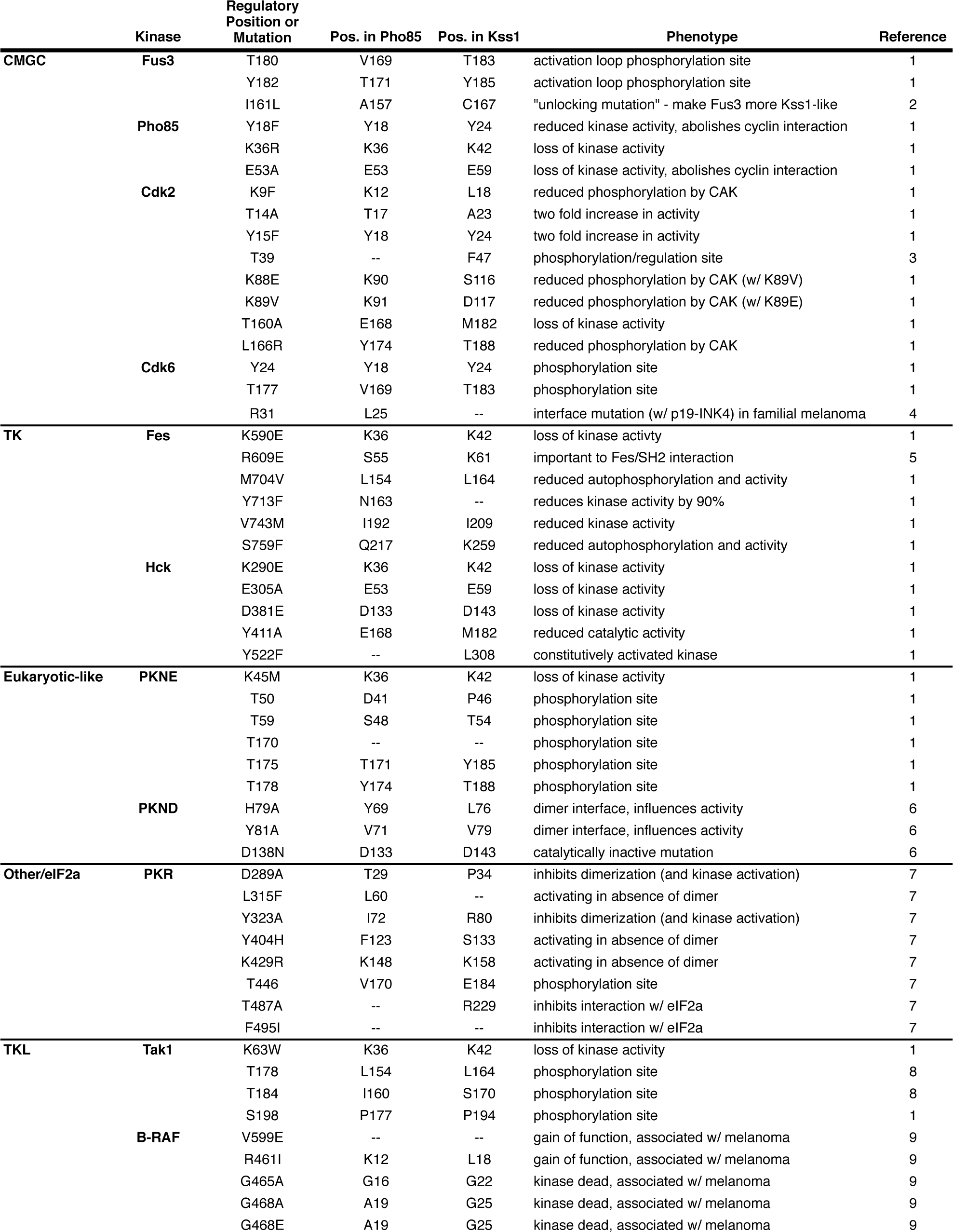
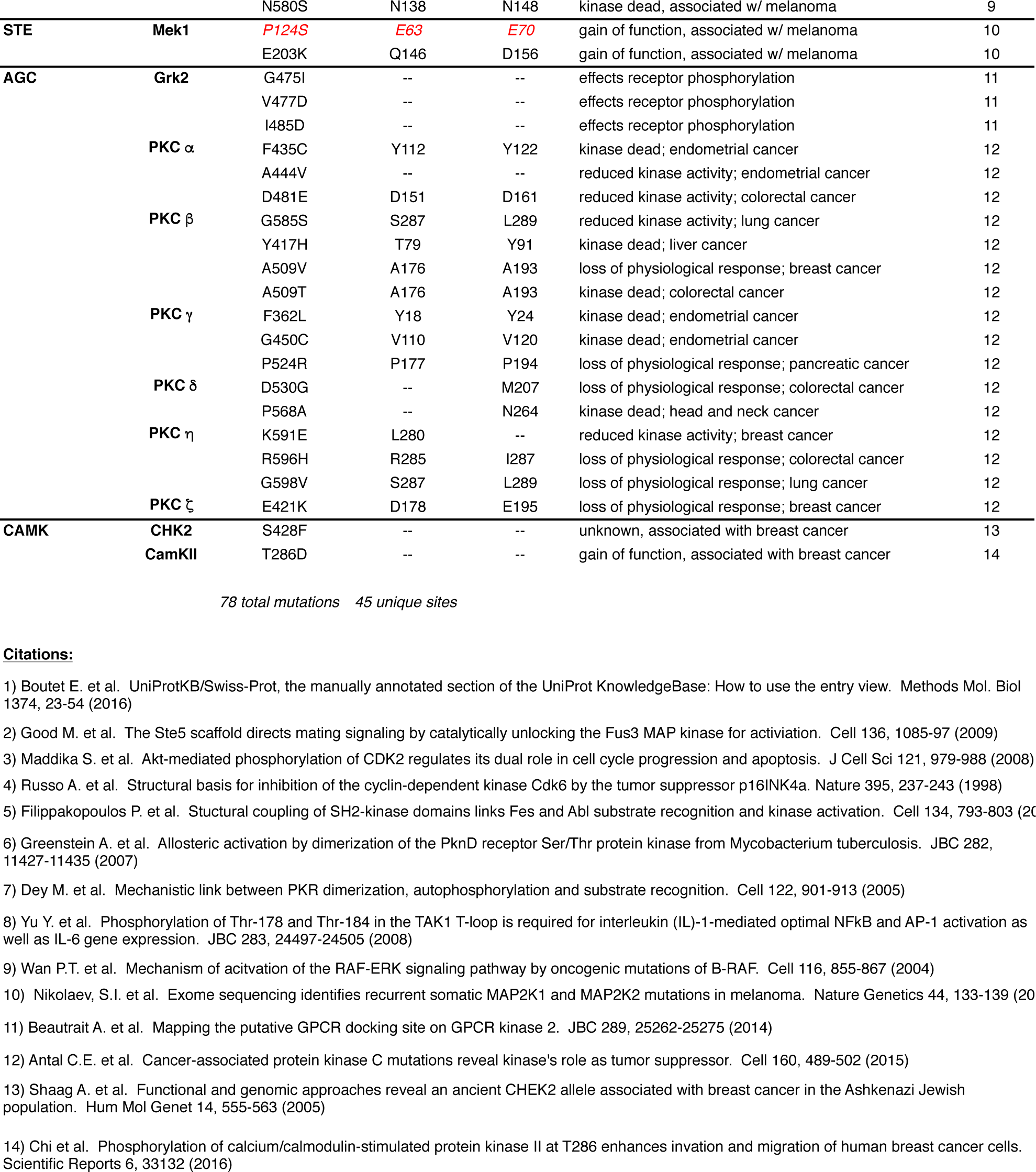
Functional Kinase Mutations (sampled across the kinome)

**Supplementary Table 8.** Statistical association between the sector, conservation and functional mutations sampled across a diversity of kinases.

**Sector positions**
| sector cutoff: | 0.95 | 0.96 | 0.97 | 0.98 |
| --- | --- | --- | --- | --- |
| CMGC alignment | 2.50E-02 | 8.65E-03 | 3.06E-02 | 1.05E-02 |
| EPK alignment | 6.82E-04 | 3.41E-04 | 7.67E-04 | 2.79E-02 |

**Conserved positions**
| Di cutoff: | 1 | 1.15 | 1.3 | 2 |
| --- | --- | --- | --- | --- |
| CMGC alignment | 5.09E-03 | 7.79E-03 | 4.41E-02 | 4.08E-02 |
| EPK alignment | 2.69E-04 | 4.19E-04 | 1.04E-03 | 2.48E-04 |

